# Effect of flagellar beating pattern on sperm rheotaxis and boundary-dependent navigation

**DOI:** 10.1101/2020.01.20.913145

**Authors:** Meisam Zaferani, Farhad Javi, Amir Mokhtare, Alireza Abbaspourrad

## Abstract

The study of navigational mechanisms used by mammalian sperm inside a microenvironment yields better understanding of sperm locomotion during the insemination process, which aids in the design of tools for overcoming infertility. Near- and far-field hydrodynamic interactions with nearby boundaries and rheotaxis are known to be some of the steering strategies that keep sperm on the correct path toward the egg. However, it is not known how the beating patterns of sperm may influence these navigational strategies. In this study, we investigate the effect of flagellar beating pattern on navigation of sperm cells both theoretically and experimentally using a two-step approach. We first isolate bovine sperm based on their rheotactic behavior in a zone with quiescent medium using a microfluidic system. This step ensures that the swimmers are able to navigate upstream and have motilities higher than a selected value, even though they feature various flagellar beating patterns. We then explore the flagellar beating pattern of these isolated sperm and their subsequent influence on boundary-dependent navigation. Our findings indicate that rheotaxis enables sperm to navigate upstream even in the presence of circular motion in their motility, whereas boundary-dependent navigation is more sensitive to the circular motion and selects for progressive motility. This finding may explain the clinical importance of progressive motility in semen samples for fertility, as the flow of mucus may not be sufficiently strong to orient the sperm cells throughout the process of insemination.

**Significance:** Finding the egg and moving toward it while traversing the complex structure of the female reproductive tract is necessary for mammalian sperm. Previous studies have shown how sperm use navigational steering mechanisms that are based on swimming upstream (i.e. rheotaxis) and along the boundaries of the female reproductive tract. We demonstrate that the performance of theses navigational mechanisms is associated with the primary characteristics of sperm motility. In fact, sperm rheotaxis is more sensitive to the motility and thus average velocity of sperm while navigation via rigid boundaries is more sensitive to the flagellar beating pattern and selects for symmetric beating. Our results can be expanded to other autonomous microswimmers and their subsequent navigation mechanisms.

## Introduction

For successful fertilization, sperm cells must traverse the distance between the location of semen deposition and the egg (1). During this transport, sperm cells require navigational mechanisms to find the correct direction in which to move (2). These navigational mechanisms rely on external stimuli, including chemical (3–5), thermal (6), and fluid mechanical (7–9) clues, which vary among different species. For instance, marine plants and animals (10, 11) release their gametes into the sea, where chemotactic behavior is observed (4, 11–13). Strikingly, the role of this chemical communication observed in marine invertebrate sperm (which is reminiscent of bacterial chemotaxis (14)) is uncertain in the navigation of mammalian sperm (13, 15–19). In contrast, *in vitro/vivo* evidence suggests that the navigation of mammalian sperm within the female reproductive tract depends more on fluid mechanical clues rather than other external stimuli (2, 7, 9, 20, 21).

The navigational system of mammalian sperm to find the correct path toward the fertilization site includes boundary-dependent navigation (20–22) and upstream swimming behavior, known as “rheotaxis” (8, 23, 24). Based on extensive studies carried out previously, the characteristics of rheotactic behavior are determined by the external flow in which the sperm are swimming, and a minimum shear rate is required for upstream swimming to emerge (23). However, boundary-dependent navigation that relies upon the hydrodynamic interactions of the swimmer with rigid walls, as well as self-propulsion and steric repulsion of sperm cells (25), is independent of the external flow and exists even in a quiescent medium. In fact, boundary-dependent navigation consists of a far-field hydrodynamic attraction of the sperm toward nearby walls (26–29), such as those of the female reproductive tract, followed by stably swimming along these boundaries (20, 21, 37, 22, 30–36). Berke et al. (27) proposed a dipole swimmer model that describes the attraction of sperm as a microswimmer with absolute progressive motility toward rigid boundaries at distances far enough from the boundaries (i.e., the far-field approximation). At close distances, swimming along boundaries is observed ubiquitously among microswimmers and Denissenko et al. (20) have demonstrated this boundary-following motion in the sperm using a microchannel and proposed it as a navigational mechanism.

Although sperm rheotaxis and boundary-dependent navigation have been studied, the intriguing unanswered question is how does the flagellar beating pattern affect these navigational mechanisms? Since sperm cells in a population feature both progressive and non-progressive motility, answering this question reveals the optimum motility mode required for sperm to be steered by such navigational mechanisms. Previous studies have explored sperm flagellar beating and its hydrodynamic interaction with rigid boundaries (20, 38, 39). However, such studies have not revealed the role of the beating pattern in sperm navigation comprehensively because they have not considered sperm rheotaxis. The dynamic flow of mucus within the female reproductive tract necessitates studying the effect of the flagellar beating pattern on sperm rheotaxis and boundary-dependent navigation concurrently. Therefore, controlled experimentation to quantify the effect of the beating pattern on different sperm navigational mechanisms is desirable. However, at shear rates at which upstream swimming occurs, characterizing the role of the beating pattern in the sperm locomotion and subsequent interactions is challenging as the flow overcomes all the non-upstream components of the motion. Furthermore, decoupling the contribution of the flow and the beating in the sperm motion is experimentally challenging in itself.

In this study, we investigate the navigation of sperm cells both theoretically and experimentally using a two-step approach. The first step is to isolate the sperm cells in a reservoir with quiescent medium via a rheotaxis-based method using a microfluidic system (40). This microfluidic rheotaxis-based isolation step ensures that all sperm cells navigate upstream via rheotaxis and have motilities higher than a selected threshold value, even though they feature various flagellar beating patterns. The second step is to study the tail beating of these sperm and their subsequent boundary-dependent navigation in a reservoir with quiescent medium to determine the influence of the tail beating pattern on this navigational mechanism. For this second step, we use phase contrast microscopy, cell tracking, resistive force theory (41), lubrication approximation (42), and finite element method simulation. This two-step approach bypasses the challenges associated with studying the flagellar beating pattern in simple shear flow yet enables us to investigate sperm rheotaxis and boundary-dependent navigation concurrently.

## Results and Discussion

To characterize the sperm flagellar beating patterns, we first isolated motile swimmers within a zone of a microfluidic chip filled with Tyrode albumin lactate pyruvate (TALP) medium using a rheotaxis-based method (40). For more details about this isolation technique, please see the Methods section. The movement of the motile sperm inside the microfluidic zone filled with quiescent medium can be seen in Movie S1, which shows that sperm with symmetric beating patterns move with progressive motion, whereas those with asymmetry in their beating patterns, i.e., one part of the flagellum consistently bends more significantly to one side, swim in circular paths. Interestingly, all these sperm have navigated upstream via rheotaxis even though they feature different beating patterns ranging from symmetric to asymmetric. In addition, the velocity of the average path (VAP) in the separated sample is higher than a threshold value (VAP ≥ 53.2 μm/s). In our previous study (40), we demonstrated that this minimum threshold is tunable via the injection rate. The various flagellar beating patterns and the tunable minimum VAP in the separated samples indicate that sperm rheotaxis is more sensitive to VAP and thus motility itself rather than the pattern of the flagellar beating.

The asymmetric flagellar beating pattern and pathway of these circular-swimming sperm were captured using phase-contrast microscopy, as shown in Fig. 1(a) and (b). Fig. 1(c) demonstrates the trajectories of nine other sperm exhibiting such circular motion, in which the centers of these circular paths move randomly over time. Initial observations also suggest the non-significant influence of the wall on the motion of the sperm with non-progressive motility compared to that of the progressive ones. In fact, due to near-wall hydrodynamic interactions, when sperm with symmetric beating patterns and thus progressive motility encounter the side wall, they begin to swim parallel along it (indicated by the blue line in Fig. 1(d)). Meanwhile, the sperm with asymmetric beating either retain their circular trajectory (indicated by the red line in Fig. 1(d)) or return to the circular trajectory after partially following the wall.

**Fig. 1.**
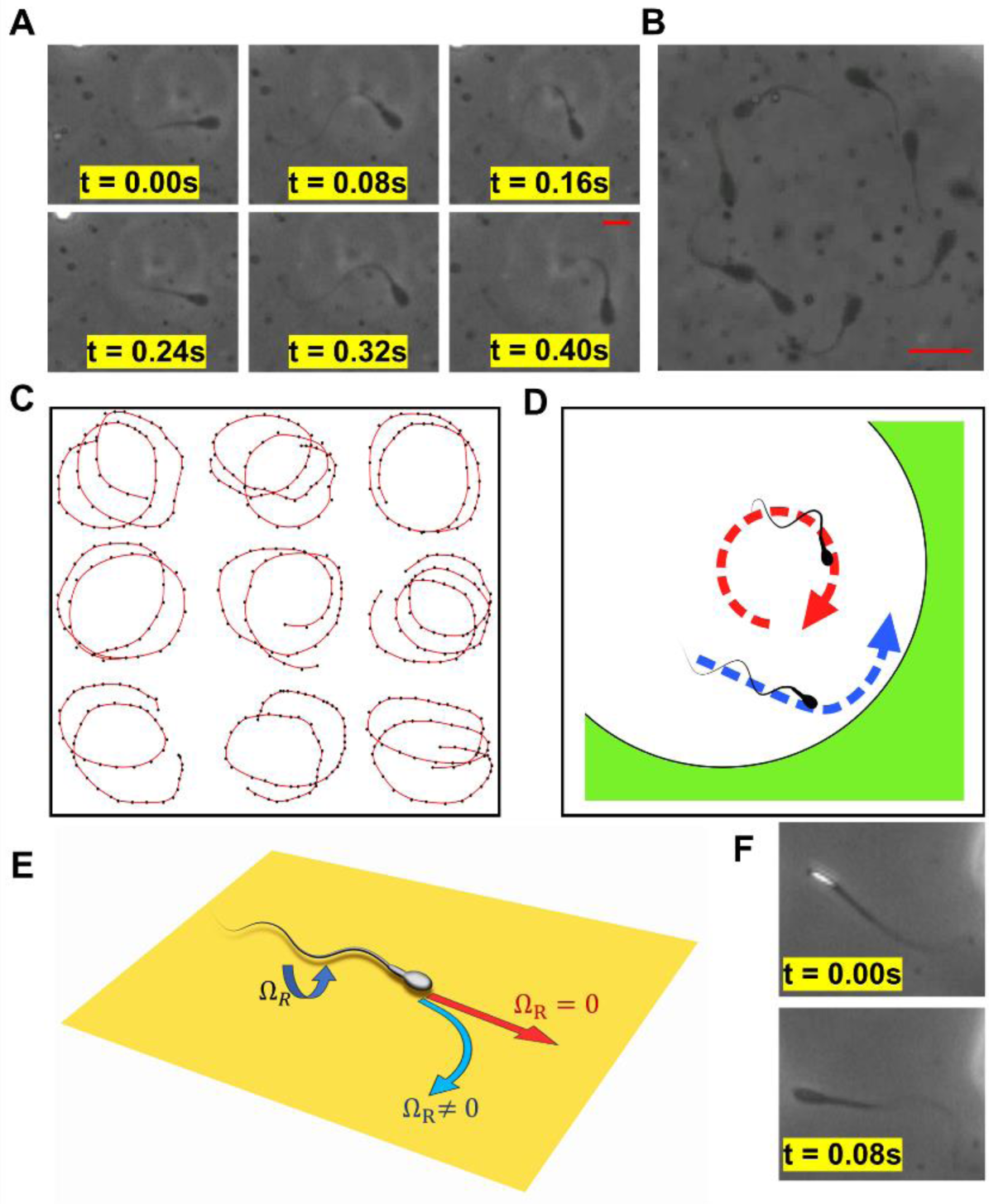
Experimental observation of sperm motion and their interactions with a curved wall. (A) The asymmetric sperm flagellar beating pattern over time, in which the thicker part of the flagellum always bends toward the left side of the sperm. (B) Image of a single sperm exhibiting intrinsic circular motion over time, acquired with phase contrast microscopy. As no twinkling was observed, it is apparent the sperm does not flip. (C) Trajectories of nine different sperm with intrinsic circular motion, in which no deterministic movement toward any direction was observed. (D) The sperm isolated within the quiescent medium feature either progressive (blue dashed line) or intrinsic circular motion (red dashed line). (E) Self-rolling motion in the swimmer results in a torque and subsequent swimming in circles. For *Ω*_*R*_ = 0, no circular motion occurs. (F) Phase contrast microscopy enables us to differentiate the sperm with and without self-rolling. Rotation in the sperm head can be identified by it twinkling. Scale bar: 10 µm

To quantitatively describe the asymmetric beating pattern, circular motion, and hydrodynamic interactions of the sperm, we first quantified the correlation between the asymmetry in the beating and the circular motion. It has been observed that in a quiescent medium, sperm (similar to *E. coli*) swim in proximity to surfaces and in large circles so that the rotation axis is perpendicular to the surface even if the motion produced by the flagellum is progressive. Tung et al. (23) and Lauga et al. (43) attributed this circular motion to the swimmer’s hydrodynamic interactions with the surface, as the head and tail of bacteria rotate counter clockwise and clockwise (or vice versa), respectively (44–46), while sperm exhibit self-rolling motion. This self-rolling motion consists of the sperm head and tail rotating in the x-z plane with Ω_*R*_, as indicated in Fig. 1(E). This rotation in the sperm head and tail generates a subsequent angular velocity that leads to the microswimmer swimming in circles in the x-y plane. Otherwise, in the absence of the self-rolling motion, a progressive sperm cell does not experience torque and swims progressively. We used phase-contrast microscopy (47) to examine whether the sperm in our experiment exhibit this self-rolling motion, in which the paddle-shaped head of the bovine sperm twinkles as it flips as demonstrated in Fig. 1(F). As can be seen in Fig. 1(b) and (c), no twinkling was observed in the sperm with intrinsic circular motion. Therefore, the hydrodynamic interactions with the top (or bottom) surface of the microfluidic chip are not responsible for the circular motion we observe, indicating the circular motion is solely due to intrinsic asymmetry in the flagellar beating pattern.

Asymmetry in the flagellar beating pattern of sperm is shown to be reliant on the intracellular Ca^2+^ concentration when it is exposed to a chemical stimulant (48, 49). For instance, the biological pathway of asymmetric beating for some marine invertebrates in a gradient of chemical stimulant, including *Ciona* and sea urchin, is known to involve Ca^2+^-sensitive proteins called “calaxin” (10) and “calmodulin” (48, 50, 51), respectively. The increase in the intracellular concentration of Ca^2+^ leads to calaxin (or calmodulin) suppressing the movement of the outer dynein arm. This suppression of the outer dynein arm then leads to asymmetric beating. Although the role of calmodulin involved in the motility of stimulated sperm is known to be central for most mammals (52) (e.g., bovine sperm (49)), the biological details of asymmetric beating at different stages of sperm motility, including activated (i.e., unstimulated) and hyperactivated (i.e., stimulated) is still unknown. Therefore, further molecular insight into asymmetric beating in mammalian sperm is required to propose a mathematical model at the single cell level that captures the molecular details of the process. Nevertheless, several studies have been carried out to propose potential mechanisms for in-plane sperm circular motion caused by asymmetric flagellar beating for both unstimulated and stimulated sperm. A well-established method to connect the asymmetry in the beating pattern of unstimulated sperm to its circular movement is to measure the intrinsic flagellar curvature (41), so that a non-zero curvature in one beat yields a circular motion. This non-zero curvature can be due to the compression of the sperm flagellum by the internal forces in a viscous fluid, which results in a buckling behavior so that symmetry in the beating breaks and thus the sperm swims in circular trajectories, as proposed by Gadelha et al. (53). Recently, Saggiorato et al. (39) studied the motion of a progesterone-stimulated tethered human sperm and proposed that a phase difference between the first and second harmonic yields a non-zero torque and thus can lead to a circular motion of stimulated sperm. Although this study brings new insights about the role of temporal harmonics in the beating pattern and its relation to the circular motion of stimulated tethered human sperm, it is not necessarily valid for the untethered, unstimulated bovine sperm in our study. Therefore, other potential mechanisms at the single cell level by which asymmetric flagellar beating of untethered and unstimulated bovine sperm leads to circular movement are required.

To determine the relation between asymmetric beating and circular motion of sperm, we first reconstructed the beating patterns of the flagellum over one beat. We then applied the Fourier transform to the beating pattern to yield its temporal frequencies (Supplementary Information Part I). Our results indicate that asymmetry in the beating pattern is associated with an increase of the main frequency (i.e., first harmonic) as well as decrease in its amplitude. Moreover, the zeroth and higher harmonics simultaneously appear in the frequency domain as the asymmetry occurs. We then described the beating pattern in the Fourier series ansatz, which includes an offset term as well as the first and higher harmonics, as described in Eq. 1,

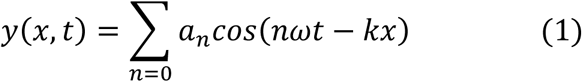

in which 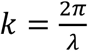 (with *λ* ≈ *L*) is the wave number, *ω* is the main frequency, and *a*_*n*_ is the amplitude of the n^th^ harmonic, including a non-thermal white Gaussian noise (54). That is, the amplitude can be described by *a*_*n*_ = *ã*_*n*_(1 + *η*_*n*_(*t*)) with ⟨*η*_*n*_(*t*)⟩ = 0 and ⟨*η*_*n*_(*t*) · *η*_*n*′_(*t*′)⟩ = *D*_*n*_*δ*_*nn*′_*δ*(*t* − *t*′). This non-thermal noise may stem from asynchrony in the collective dynamic (55–57) of the dynein motor proteins that are responsible for transport along microtubules within the sperm flagella (3, 10).

Applying resistive force theory on the wave function described by Eq. 1, we can calculate the forces produced by each segment of the flagellum in the tangential (x) and normal (y) directions using 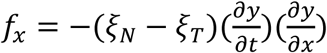 and 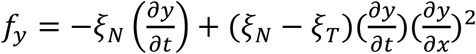, in which *ξ*_*N*_ and *ξ*_*T*_ are drag coefficients in the normal and tangential directions. Therefore, the time-averaged forces produced by each segment of the flagellum in the tangential 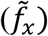 and normal 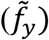 directions can be described by:

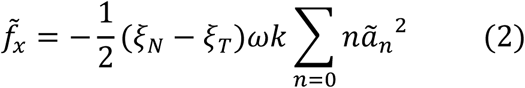

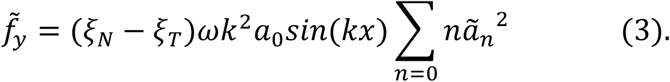

Integrating 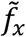 and 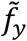 over the flagellum, the total forces produced in the tangential and normal directions are 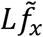 and zero, respectively. Although the emergence of the zeroth harmonic does not lead to a net normal force and subsequent translational motion, it produces a torque

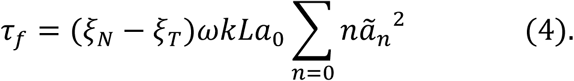

The obtained relations for the produced force and torque by the flagellum reveal that while the tangential force is correlated to the characteristics of the first and higher harmonics, the amplitude of the zeroth harmonic is involved in the torque as well. Applying the zero net torque and force constraint, the tangential and angular velocity are found to be correlated through *a*_0_:

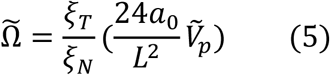

in which 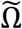 is the average angular velocity (Supplementary Information Part II).

To validate the analysis, we experimentally measured the angle sperm sweeps out in one beat (Δ*θ*), in which Δ*θ* is the difference between the deviation of the sperm head direction from the average path to its left (*θ*_1_) and right (*θ*_2_) sides (Fig. 2A). Fig. 2B displays the measured Δ*θ* values for sperm with progressive motility and circular motion. As can be seen, the average of Δ*θ*(*t*) (i.e., 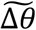) for sperm with circular movement are significantly higher than that of sperm with progressive motility. Considering that 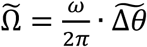, we measured the normalized angular velocity 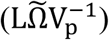 and curvature of the sperm path 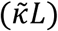 with regards to the amplitude of the zeroth harmonic, as shown in Fig. 2C. Based on Fig. 2C, we determined 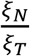 to be 1.93 ± 0.33, which is comparable to previously reported values (58). Based on these experimental results and the agreement they show to the relations we derived from resistive force theory, we conclude that the circular motion in the sperm motility is attributed to the zeroth harmonic in the flagellar beating.

**Fig. 2.**
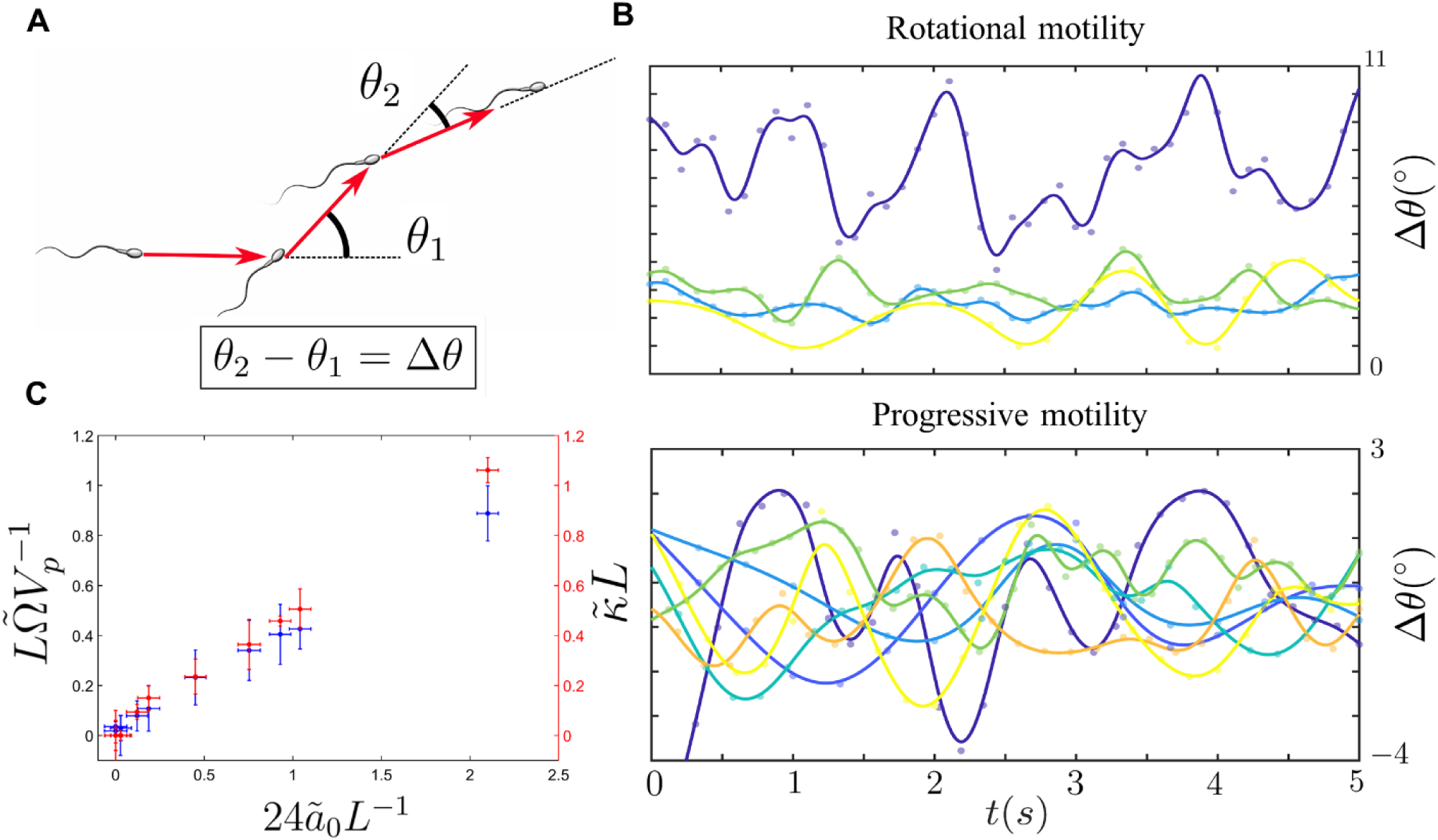
Intrinsic circular motion caused by the presence of the zeroth harmonic. (A) Schematic of sperm motion featuring intrinsic circular motion. *θ*_1_and *θ*_2_ are the deviation in the sperm direction towards the left and right of the mean path. (B) Δ*θ* measured for sperm with circular motion (top) and progressive motility (bottom). The average value of Δ*θ* is significantly higher in sperm with circular motion. (C) The normalized angular velocity (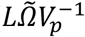, blue) and curvature of the sperm path (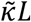, red) plotted versus the amplitude of the zeroth harmonic 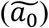 to experimentally confirm the results obtained from resistive force theory. 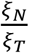 was determined to be 1.80 ± 0.34 and 2.07 ± 0.31 for the blue and red data points, respectively. The average of these values reported as 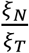 in the main text.

We also demonstrate the effect of non-thermal Gaussian noise (33) in the amplitude of the zeroth harmonic on the sperm trajectory. Based on the results gained from the resistive force theory analysis, the noise in the zeroth harmonic yields a similar noise in the angular velocity. Therefore, the angular velocity can be described as 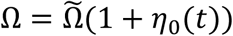. Including the non-thermal white Gaussian noise in the angular velocity and simulating the sperm motion at different signal-to-noise ratios (SNR), we found that this noise leads to stochastic movement of the center of the sperm’s circular path, as can be seen in Fig. 3A, which is consistent with our experimental observation of bull sperm movement shown in Fig. 1C. These findings suggest that the inconsistency in the sperm’s circular path is a consequence of the noise in the amplitude of the zeroth harmonic. In addition, we found the sperm are more capable of maintaining their circular path at higher SNR values (SNR = 50, 55, 60), and thus we observe less movement at the center of these paths, as can be seen in Fig. 3A. At lower SNR values, the sperm motion contains more stochasticity and thus covers a larger domain. We repeated the simulation for 1000 sperm cells to find the diffusivity of the circular path’s center (i.e., *D*_*c*_) at different SNR values, and as can be seen in Fig. 3B, *D*_*c*_ is inversely correlated to the SNR. Accordingly, we expect the distance of the circular path’s center from its initial location, *r*_*c*_(*t*) to increase over time with (*D*_*c*_*t*)^0.5^. Fig. 3C shows the *r*_*c*_(*t*) obtained at different SNR values, which confirms the localization of the sperm at high SNR and the increase of *r*_*c*_(*t*) in time with *t*^0.5^.

**Fig. 3.**
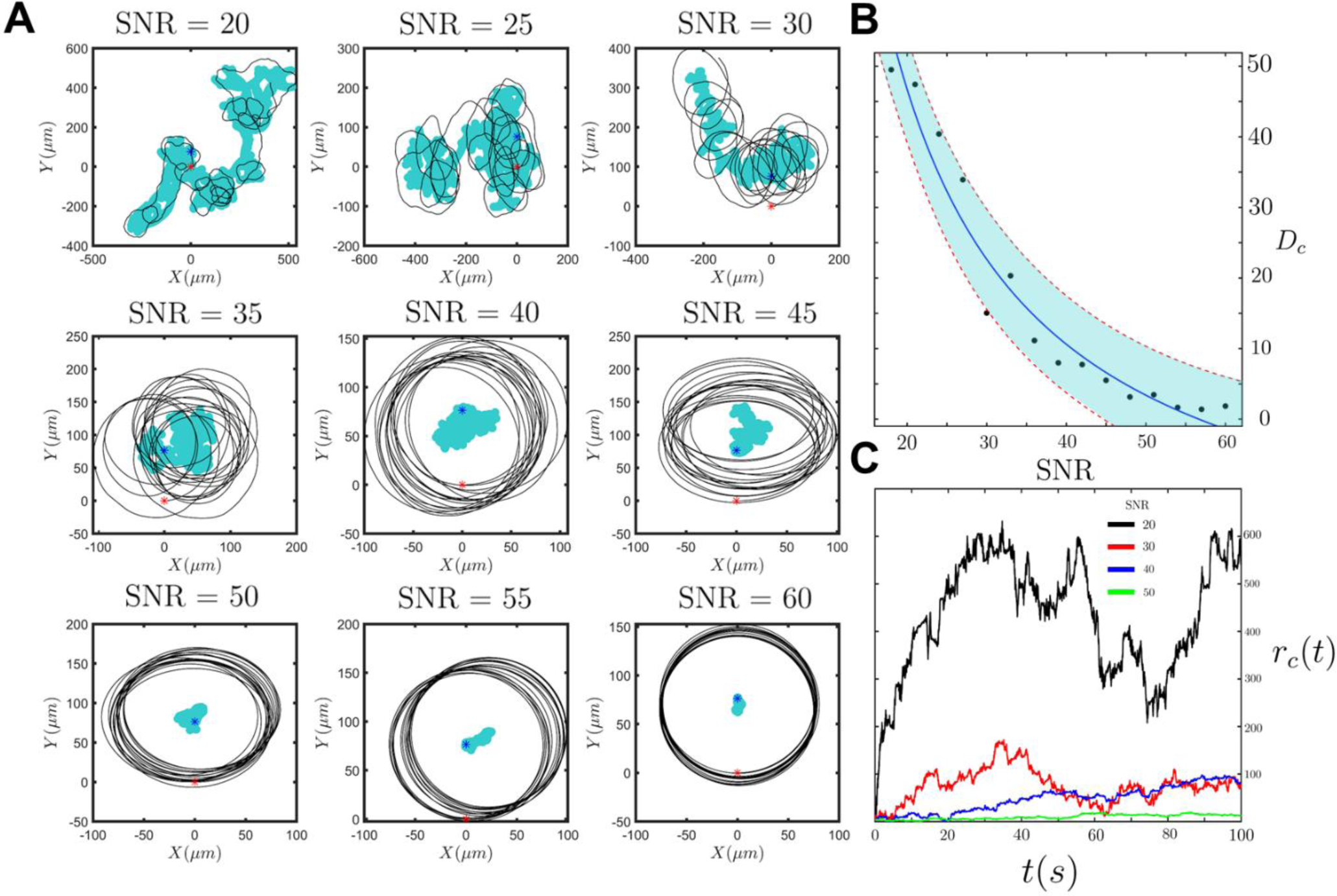
Noise in the angular velocity and the corresponding diffusivity of the circular path’s center. (A) The sperm trajectory at 9 different signal-to-noise ratios (20 ≤ *SNR* ≤ 60). Higher SNR values yield more consistent and deterministic circular motion. (B) The diffusivity of the circular path’s center vs. SNR. Each point in the figure was obtained by simulating the motion of 1000 sperm cells with similar initial conditions. (C) The distance of the circular path’s center from its initial location with respect to time. The obtained results are at 9 different SNR; 4 of which are demonstrated in this figure.

Berke et al. (27) modeled the far-field approximation of the flow generated by a microswimmer with progressive motion as a pusher force dipole. Therefore, we used the sperm dipole model, including the components of the circular motion, to calculate the sperm contribution to the fluid flow and thus the hydrodynamic interactions inside the quiescent zone at distances adequately away from the curved sidewall (> 2 × sperm length). The proposed model is depicted in Fig. 4A, in which *f* is the tangential force produced by the sperm flagellum, *f*′′ is the perpendicular force corresponding to the torque caused by asymmetric beating, and *f*′ is the drag force required for the torque-free condition (26). The magnitude of *f*′ and *f*′′are equivalent to each other while the 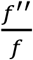 ratio (i.e., γ) is equal to 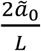 (see Eq. S38). The contribution of the sperm’s active swimming to the fluid flow is the solution of the Stokes equation for the proposed model shown in Fig. 4A. The general form of the Stokes equation for such a swimmer is

**Fig. 4.**
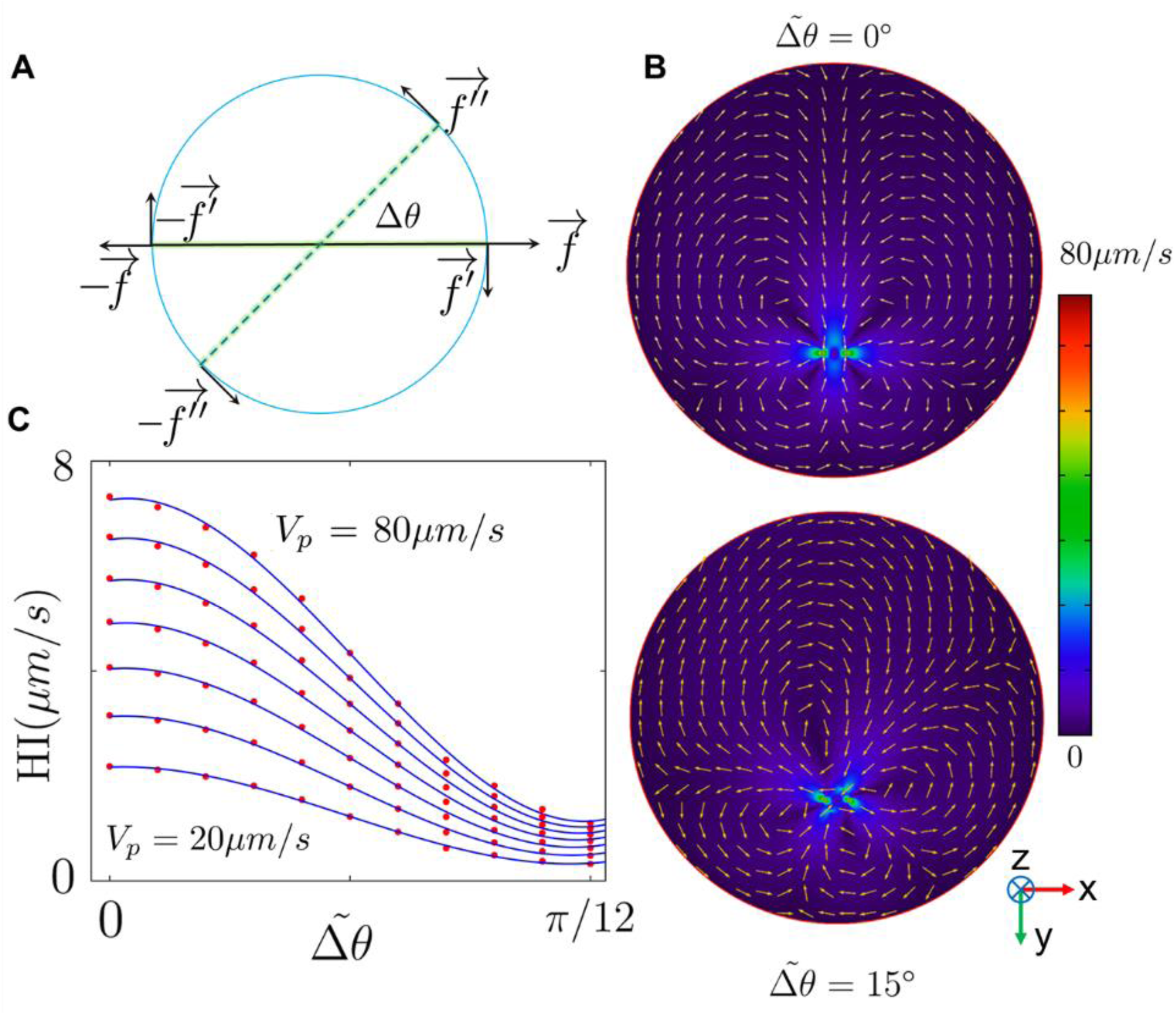
Dipole swimmer model and far-field hydrodynamic interactions. (A) The dipole swimmer model, including the components of the circular motion (*f*′ and *f*′′). (B) The velocity field obtained from solving the Stokes equation for the dipole swimmer model in a quiescent medium. The colors represent the magnitude and the arrows are normalized to visualize the direction of the vector field. (C) The attraction caused by the presence of the wall decays as we increase Δ*θ*.

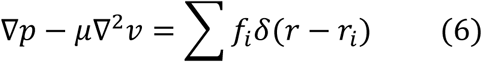

in which *p* is the pressure, *μ* is the dynamic viscosity of the TALP medium (0.94 mPa/s), *v* is the fluid velocity, *r* is the position, *r*_*i*_ is the position of the point force *f*_*i*_, and *δ* is the Dirac delta function. To solve the Stokes equation, we can either use Green’s function, known as the “Stokeslet” description (42, 59), or solve the equation numerically. The analytical expression for the swimmer model using the Stokeslet description and mirror image (46, 60) method is explained in Part IV of the Supplementary Information.

The near- and far-field flow produced by sperm has been studied previously (61). However, for the sake of simplicity and precision, we carried out finite element method simulations in a cylindrical domain similar to the quiescent zone to numerically solve the Stokes equation along with mass conservation (62). Considering the sperm swim in a quasi-2D plane that is located ∼5 μm below the top surface and parallel to it (63), we obtained the velocity field imposed by the flagellar beating for 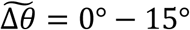 (Fig. 4B). We then integrated the net flow in the y direction (Fig. 4B) imposed on the sperm body, which is caused by the presence of the no-slip walls, to calculate the hydrodynamic interaction (i.e., HI), the results of which are demonstrated in Fig. 4C. The velocity field generated by the progressive flagellar beating 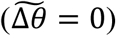 in the presence of the no-slip boundaries leads to the attractive hydrodynamic interactions (> 0), as indicated in Fig. 4C, which agrees with previous studies (22). As we added and increased the components of the circular motion, the attractive hydrodynamic interactions decayed for a constant progressive velocity of the sperm (20 *μm*/*s* <*V*_*p*_ < 80 *μm*/*s*), indicating that the motion of the microswimmer is less influenced by nearby boundaries as components of circular motion emerge in the motion. To gain a better understanding about the mechanism of this reduction in hydrodynamic attraction, we simulated the flow field produced by components of the progressive 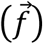 and the circular motion (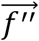 and 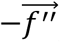) and their corresponding drags (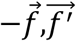 and 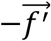) separately (Fig. S4). The flow field produced by the progressive component and its corresponding drag is outward in the front and rear of the swimmer, while inward on the sperm’s right and left sides. In contrast, the flow field produced by the components of the circular motion and their corresponding drags are inward in the front and rear of the swimmer, while outward on the sperm’s right and left sides. Accordingly, as the components of the circular motion becomes larger, more of the flow imposed by the progressive motion is damped by that of the circular motion and therefore the hydrodynamic attraction decreases with sperm circular motion.

This decay in the far-field hydrodynamic interaction is also predicted by the analytical expression for the swimmer model using the Stokeslet description. In fact, the attraction of the sperm toward the wall in the presence of the components of the circular motion can be described by:

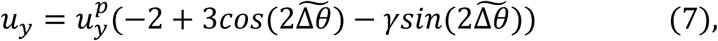

in which *u*_*y*_ and 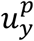 describe the far-field attraction with and without the components of the circular motion. Neglecting *γ*, an increase in 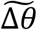 results in a decrease of attraction to the walls. Interestingly, at 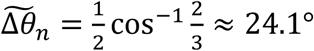 and corresponding curvature of 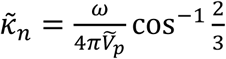, the swimmer experiences no attraction toward the walls and becomes neutral.

At the near-wall condition (< the sperm body length), the dipole approximation is no longer valid (35) and the sperm-wall interaction can be understood by near-field approximations, as previous theoretical (38, 64) and simulation-based (65) studies suggest. We categorize the sperm interaction with the curved sidewall into four different types: 1) a progressive sperm encounters the wall, rotates, and follows it, as can be seen in Fig. 5A(I); 2) a non-progressive sperm encounters the wall, follows it temporarily, and detaches (Fig. 5A(II)); 3) a non-progressive sperm that does not contact the wall (Fig. 5A(III)); and 4) a non-progressive sperm that encounters the wall and stays still or moves slowly along it (Fig. 5A(IV)). These categories can be seen in Movie S2. To interpret these near-wall interactions, we first used surface contact force analysis, in which the wall influence on the sperm movement is modeled as a normal force.

**Fig. 5.**
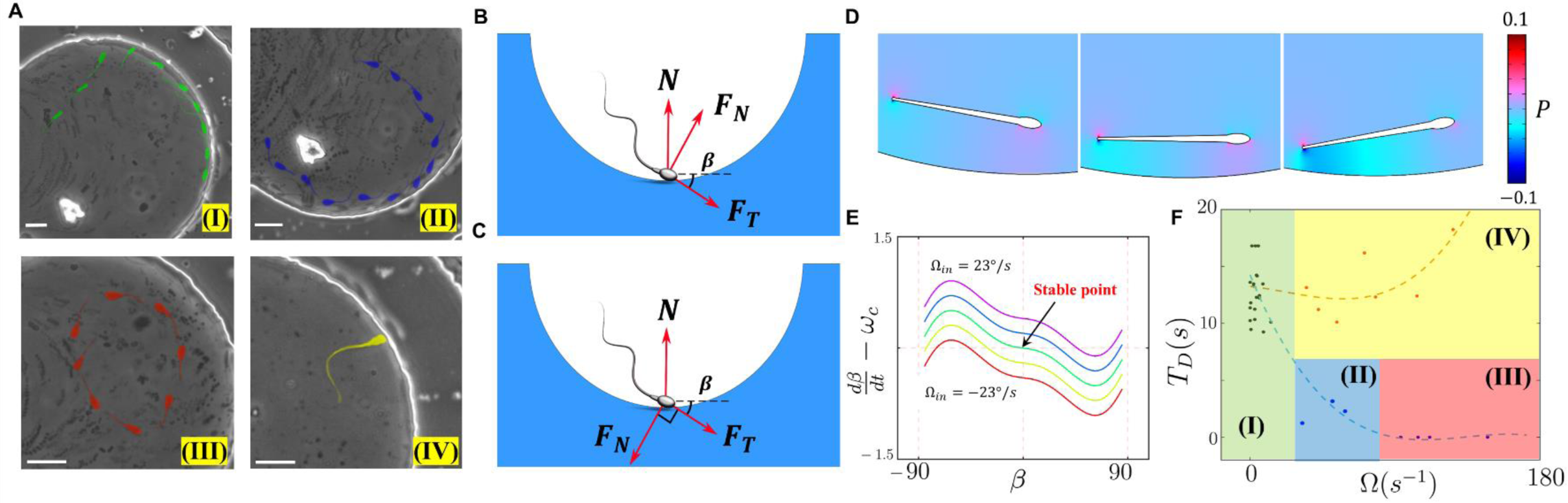
Near-field interactions and the susceptibility of the sperm to follow the wall. (A) Sperm with (I) progressive motility that is susceptible to the wall, (II) low intrinsic circular motion and partial susceptibility to the wall, and (III) high intrinsic circular motion and zero susceptibility to the wall. A non-progressive sperm that encounters the wall and stays still is also depicted in (IV). Scale bar: 10*μm* (B) The forces produced by the sperm flagellum and the normal surface contact force (*N*). This arrangement describes Fig. A(I) and (II) and (III). (C) The forces produced by the sperm flagellum and the normal contact force. This arrangement describes Fig. A(IV). (D) The pressure distribution caused by sliding of sperm nearby the wall at *β* = −20°, 0°, and 20°, from left to right. This pressure distribution is used to calculate the torque and angular velocity imposed on the sperm by the presence of the wall. (E) Dynamic of the sperm incidence angle in the phase-plane, depicted at different intrinsic angular velocity values (−23 °/*s* ≤ Ω_*in*_ ≤ 23 °/*s*). (F) The sperm detention time on the curved wall measured for sperm with different intrinsic angular velocities. Sperm with |Ω_*in*_| > 76°/*s* (red box) do not contact the wall, while the ones with 8° ≤ |Ω_*in*_| ≤ 76° (blue box) are partially susceptible to the wall and follow it temporarily. Sperm with |Ω_*in*_| ≤ 8° (green box) are capable of following the wall without detaching. The detention time for sperm cells corresponding to (IV) in Fig. 5(A) and Fig. 5(C) are presented in the yellow box.

When sperm is in contact with the wall surface there is a positive surface contact force, whereas a surface force of zero corresponds to detachment of the swimmer from the wall (66). Consider a sperm that encounters the wall of the quiescent zone with an incident angle of β, as depicted in Fig. 5B. Under a zero-net force constraint, the normal surface force becomes *N* = *F*_*T*_*sin*(*β*) − *F*_*N*_*cos*(*β*), where *F*_*T*_ and *F*_*N*_ are the tangential and perpendicular forces, respectively. The threshold angle that corresponds to the *N* = 0 situation (i.e., *β*_*th*_) is equal to tan^−1^ *γ*. For *β* < *β*_*th*_, the normal surface force becomes negative and no contact occurs accordingly. Hence, the sperm does not follow the wall. Since an increase in *γ* leads to higher *β*_*th*_, sperm with greater *γ* values are less likely to contact and follow the wall.

For incident angles greater than *β*_*th*_ (where the sperm-wall contact occurs), an increase in *F*_*N*_ leads to a smaller N, and thus easier detachment from the wall results. For the other condition, in which the direction of the perpendicular force is opposite (Fig. 5C), the surface force becomes greater (*N* = *F*_*T*_*sin*(*β*) + *F*_*N*_*cos*(*β*)) and detachment is more difficult. However, since the final orientation of the swimmer at the contact point is tilted, the component of the perpendicular force along the wall overcomes the tangential force, and subsequently the sperm movement along the wall becomes slower. At the extreme condition, the sperm cell stands still.

Although surface contact force analysis can be used to gain a general notion of near-field interactions, its only valid under circumstances where contact occurs. Therefore, we also developed a hydrodynamic explanation for near-field interactions to obtain more quantitative characterization. We used lubrication theory as a platform, where the sperm distance from the wall was assumed to be much smaller than its length (42). We then solved the Stokes equation and extracted the pressure distribution for sperm (Fig. 5D) at different incident angles (−90° < *β* < 90°) and constant progressive velocity (*V*_*p*_ = 80 *μm*/*s*). Given that the contribution of pressure in the stress tensor dominates that of the viscous stress (42) (*pI* ≫ *μ*(∇*v* + ∇*v*^*T*^)), the torque exerted by the wall was calculated and the corresponding angular velocity is

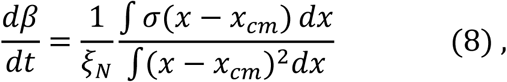

in which *σ* is the stress tensor and *x*_*cm*_ is the coordinate of the sperm’s center of mass. The obtained angular velocity as a function of *β*, including the effect of Ω_*in*_, is shown in Fig. 5E. As can be seen in Fig. 5E, for Ω_*in*_ = 0, the stable point occurs at *β* = 0 with 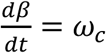, which is the required angular velocity to follow the boundary of the curved wall (Supplementary Information part IV). Whereas for Ω_in_ > 0, which corresponds to the configuration of Fig. 5B, 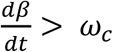 at *β* = 0, meaning it is not stable when following the boundary and the sperm subsequently detaches from the wall. For Ω_*in*_ < 0 (Fig. 5C), 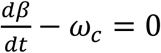 at *β* < 0, which results in the tilted orientation of the sperm at its contact point, and thus harder detachment and slower boundary-following motion.

To confirm our results obtained from the surface force analysis and lubrication approximation, we experimentally measured the time of sperm detention on the wall (i.e., *T*_*D*_) for different Ω_*in*_ (Fig. 5F). The experimental results for *T*_*D*_ indicate that the sperm exhibiting intrinsic circular motion either do not contact the wall (|Ω_*in*_| > 76°/*s*) or swim on the wall for shorter times (8 °/*s* < |Ω_*in*_| < 76°/*s*) compared to progressive sperm, which is consistent with the findings of the surface force analysis. The detention time for sperm cells corresponding to (IV) in Fig. 5(A) and Fig. 5(C) are presented in the yellow box in Fig. 5(F), which confirms that the components of the circular motion in sperm can lead to higher detention times and harder detachments, as we anticipated. Based on these experimental data, and similar to the far-field case, sperm with progressive motility are the most susceptible to near-field boundary-dependent navigation.

By isolating sperm cells based on their rheotactic behavior using a microfluidic system, we demonstrated that sperm cells are able to navigate via upstream swimming while they feature a continuum of asymmetry in their flagellar beating patterns. To identify the role of asymmetric beating on the boundary-dependent navigation, and then compare it to rheotaxis, we then investigated sperm motion after isolation in a quiescent zone with curved walls. Our results indicate that asymmetric flagellar beating patterns cause sperm to swim in circles. This circular motion of the sperm the far-field hydrodynamic interactions so that sperm with intrinsic circular motion are less attracted to walls. Likewise, at distances closer to the wall, the sperm with non-progressive motility are less likely to navigate along the wall in comparison to completely progressive sperm.

Based on these results, we conclude that boundary-dependent navigation is more sensitive to the beating pattern compared to sperm rheotaxis; whereas sperm rheotaxis is more sensitive to the motility (VAP). Accordingly, the findings of this paper, accompanied with the clinical correlation between fertility and progressive motility in sperm samples (67), suggests that at some points during the fertilization process, boundary-dependent navigation plays a central role. The findings of this study provide a comprehensive understanding of sperm locomotion during the fertilization process before reaching the fertilization site and thus can be used to improve the conventional tools for infertility diagnosis and treatment (68).

## Materials and Methods

### Semen sample preparation

Commercially available cryopreserved bovine semen samples were kindly provided by Genex Cooperative (Ithaca, NY) in milk and egg yolk-based extender in plastic straws. The semen in the straws were thawed in a 37 °C water bath and then diluted 1:3 with TALP (the sperm culture medium). After dilution, the viscosity of the samples was ∼2.1 mPa s^-1^. The initial sperm concentration in the semen samples were ∼100 million/mL and after diluting with TALP, reduced to a quarter of the initial concentration. The motility of the semen sample after dilution ranged from 20–30%. We used 10 different semen samples in both milk and egg yolk-based extender. The TALP recipe was as follows: NaCl (110 mM), KCl (2.68 mM), NaH_2_PO_4_ (0.36 mM), NaHCO_3_ (25 mM), MgCl_2_ (0.49 mM),CaCl_2_ (2.4 mM), Hepes buffer (25 mM), glucose (5.56 mM), pyruvic acid (1.0 mM), penicillin G (0.006% or 3 mg/500 ml), and bovine serum albumin (20 mg/ml).

### Micro fabrication process and semen injection

The microfluidic device was made of polydimethylsiloxane using a standard soft lithography protocol (69, 70). The diameter of the curved quiescent zone was 200 µm and the height of the chamber was 25 µm. The diluted semen was injected into the microfluidic device using gravitation and the flow generated in the channel was controlled by changing the height of the semen container. Since sperm rheotaxis emerges under very low shear rate (0.6 s^-1^), using gravitation instead of conventional syringe pumps is a more efficient way to obtain and control low flow rates.

### Rheotaxis-based sperm isolation and phase-contrast microscopy

To isolate motile bovine sperm inside the quiescent zone, we utilized a microfluidic corral system(40) that isolates motile swimmers based on their ability to move upstream. As we injected the sample with an injection rate of 1.2 mL/h, sperm with motilities higher than 53.2 μm/s can swim upstream and enter the quiescent zone (which is filled with TALP), allowing us to study the sperm movement with minimal fluid mechanical noise. The movement of the sperm cells were acquired with a phase-contrast microscope, where flipping of the sperm head leads to subsequent phase shifts in the light passing through, leading to variation in the observed brightness (i.e., a twinkling effect).

### Cell tracking and zeroth harmonic measurement

The sperm trajectories and other motility related characteristics were analyzed with ImageJ and a custom MATLAB code. To measure the amplitude of the zeroth harmonic, we measured the maximum amplitude of the beating towards the left (*y*_*L*_) and right (*y*_*R*_) sides of the swimmer, so that the magnitude of the zeroth harmonic is:

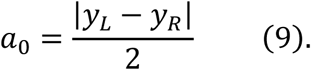

The error bars in Fig. 2(C) result from measuring *a*_0_ at 10 different beats, while the error bars of 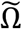 and 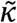 in the same figure were obtained from 10 circles of movement.

### Numerical simulation

#### Beating pattern reconstruction

To reconstruct the beating pattern of the sperm, we posited the temporal part of the flagellar beating could be described by two Sine and Cosine function defined on the interval of [*π* − *ϕ*_0_, *ϕ*_0_] so that *ϕ*_0_ ∈ [*π*, 2*π*]. Later, by evenly extending the function (71), we obtained the beating patterns so that *ϕ*_0_ determines the asymmetry in the beating; e.g. *ϕ*_0_ = 2*π* corresponds to completely symmetric beating and thus progressive motility, while *ϕ*_0_ < 2*π* results in asymmetry. We then applied a fast Fourier transform on the beating patterns to determine their temporal frequencies. These steps were performed using MATLAB (version R2017a).

#### Finite element method simulations

To obtain the velocity field imposed by the dipole swimmer model and determine the far-field hydrodynamic interactions, we first imported the structure of the quiescent zone in the COMSOL MULTIPHYSICS (version 5.2) platform. Two orthogonal Gaussian pulse functions (defined in the *x* and *y* directions) were used to define each point force in the dipole swimmer model. The mathematical form of the pulse is a 2D Gaussian distribution

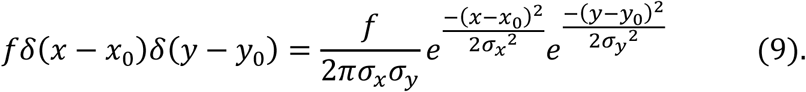

We use *x*_0_,*y*_0_ to move and *σ*_*x*_, *σ*_*y*_ to focus the point forces arbitrarily. This strategy was chosen to lower the computational costs and prevent issues related to using small volumetric forces and their associated meshing problems in the finite element method. Finally, solving the Stokes and mass conservation equations for different Δ*θ* values, we obtained the results demonstrated in Fig. 4B. Then by integrating the velocity field imposed by the sperm, we obtained the hydrodynamic interactions in the y direction, as shown in Fig. 4C.

To find the torque imposed on the sperm at near-field through the lubrication approximation, we first solved the Stokes equation for the schematic shown in Fig. 5D at different incident angles. Exporting the pressure distribution and assuming the sperm’s center of mass was located on the flagellum and twice closer to the head than to the tail, the torque imposed by the wall on the sperm and the subsequent angular velocity were calculated.

## Acknowledgement

The authors would like to thank S. H. Cheong for providing the bull semen samples and phase-contrast microscope, as well as D. L. Koch, C. K. Tung, J. Fan, and M. Esmaily for helpful discussions about hydrodynamic interactions, the dipole swimmer model, and the surface force analysis used here. This work was performed in part at the Cornell Nano Scale Facility, an NNCI member supported by NSF Grant NNCI-1542081.

## Author contributions

M.Z. and A.A. designed the research. M.Z. performed the experiments and analyzed the data. M.Z and F.J. conceived the theoretical parts. M.Z. and A.M. performed the FEM simulations. M.Z. and A.A. wrote the paper.

## Supplementary Information

### I. Reconstruction of sperm beating patterns and Fourier analysis

To model beating patterns that resemble that of the sperm flagella, we studied the pattern in one cycle of flagellar beating using a traveling sine wave with a temporal phase of *ϕ*(*t*) = *ωt* (with *ω* = 40*π* Hz as the angular frequency) in the range of *π* − *ϕ*_0_ ≤ *ϕ*(*t*) ≤ *ϕ*_0_ so that *ϕ*_0_ ∈ [*π*, 2*π*]. In turn, we constructed the even extension of the partial sine wave to form the flagellar beating function over time. Fig. S1 shows these constructed patterns, which resemble the flagellar beating of the sperm observed inside the quiescent zone. Here, the completely symmetric beating (*ϕ*_0_ = 2*π*) corresponds to sperm with absolute progressive motion, while asymmetry within the flagellum motion (*ϕ*_0_ < 2*π*) results in intrinsic circular motion. To analyze the beating patterns and the resulting sperm motion, we applied the Fourier transform to yield the temporal frequencies (Fig. S2). Interestingly, with increasing temporal asymmetry in the beating pattern, the frequency of the main (first) harmonic increases while its amplitude decreases. Moreover, the zeroth and second harmonics simultaneously appear in the frequency domain.

**Fig. S1.**
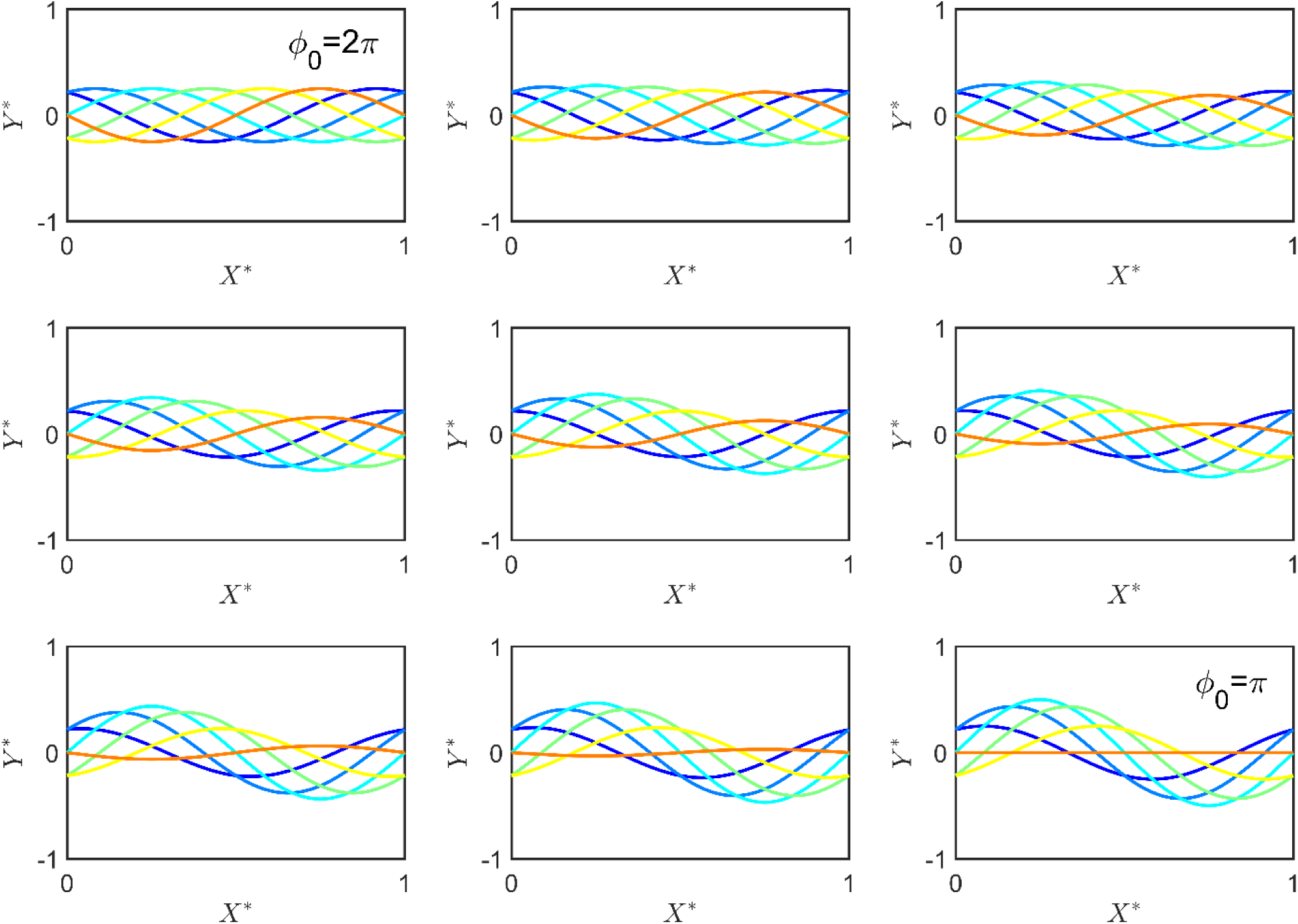
The beating pattern of the sperm flagellum. The *ϕ*_0_ = 2*π* situation corresponds to symmetric beating, and thus absolute progressive motility. As *ϕ*_0_ decreases, asymmetry in the beating emerges and the motility includes a circular motion. *x*\* and *y*\* are normalized *x* and *y* axes as the sperm motion can be described in 2D Cartesian system.

**Fig. S2.**
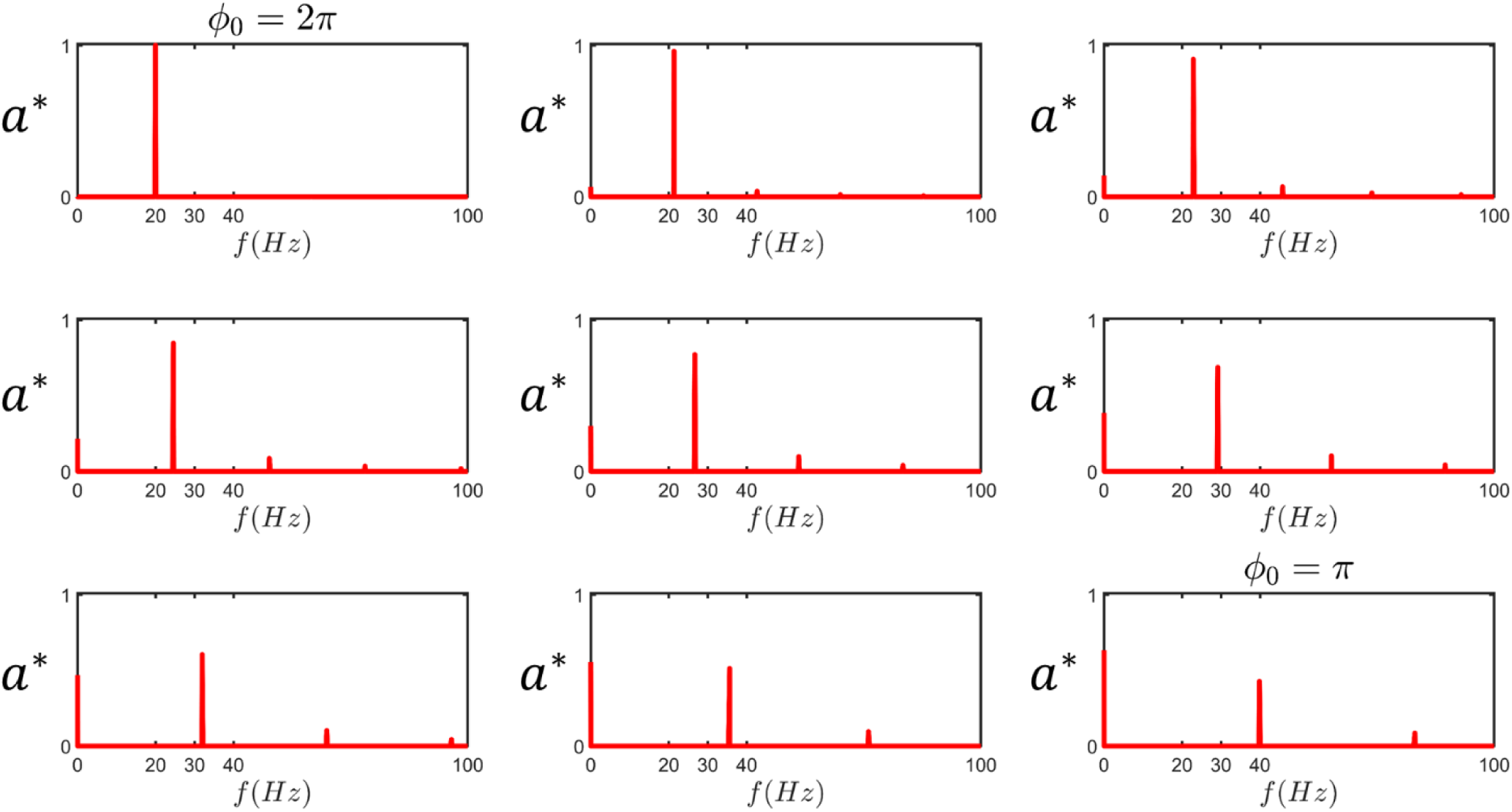
The harmonics of the sperm beating. At *ϕ*_0_ = 2*π*, the beating pattern comprises a single frequency equal to 20 Hz. As asymmetry emerges in the beating, the main frequency shifts and higher harmonics along with the zeroth harmonic appear in the spectrum. ***a***\* is the normalized amplitude of the harmonics.

### II. Resistive force theory

In this section we use resistive force theory to derive equations describing the forces produced by each segment of the flagellum and follow the presentation by Friedrich *et al*. (ref 41) and Saggiorato *et al*.(ref 39). The velocity of each segment in the y direction (*V*) can be decomposed into its tangential and normal components, *V*_*T*_ and *V*_*N*_, using *α*, which is the tangential angle (S1–4).

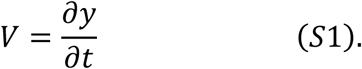

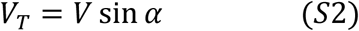

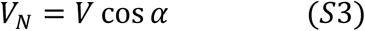

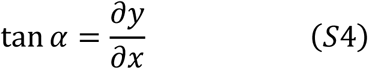

Relying upon resistive force theory, the forces produced by each element in the tangential and normal directions are linearly related to the velocity in those directions:

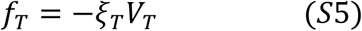

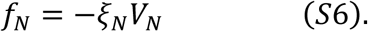

In which, *ξ*_*T*_ and *ξ*_*N*_ are drag coefficients in the tangential and normal directions. Since the amplitude of all harmonics are small in comparison to the sperm length, we can make the following assumptions:

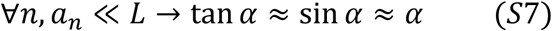

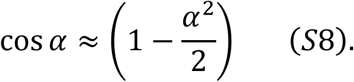

Using the approximations in Eq. S7–8, we can write out the tangential and normal velocities and forces using Eq. S9–12.

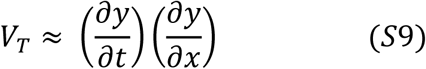

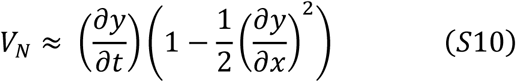

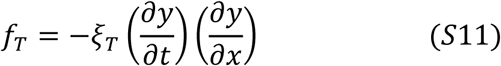

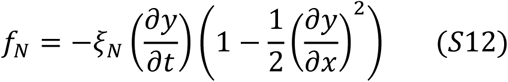

**Fig. S3.**
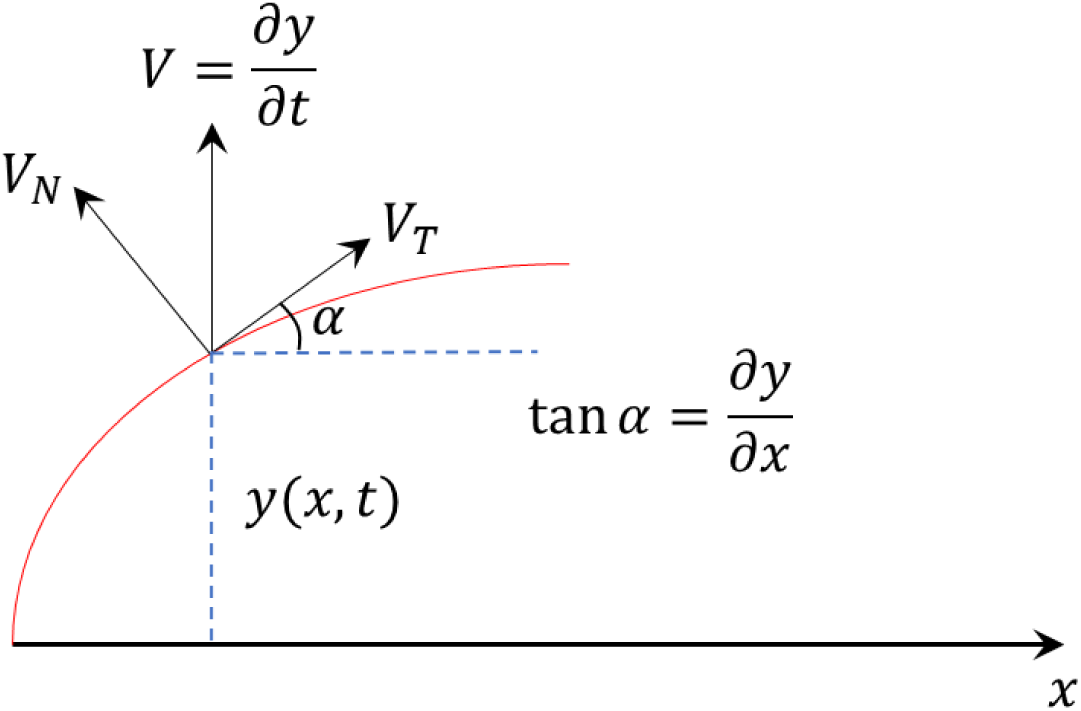
A segment of flagellum.

The force produced by each segment in the x and y directions are described by Eq. S13–14.

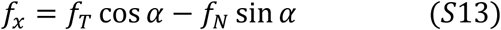

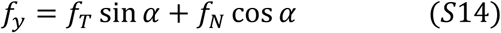

Plugging Eq. S7, 11–12 into Eq. S13–14:

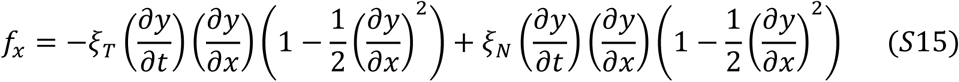

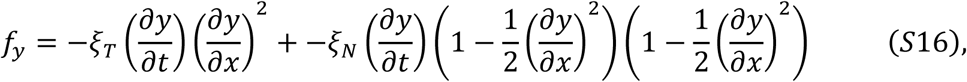

yields Eq. S17–18, which are the forces produced by the flagellum in the x and y directions.

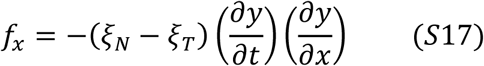

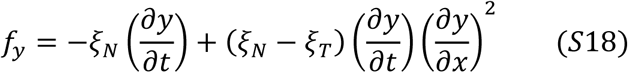

### III. Force and Torque produced by the flagellum

#### 1. Force in the x direction

The forces produced by a segment of flagellum moving with *y*(*x, t*) in the tangential and normal directions can be described by Eq. S17 and S18, where *ξ*_*T*_ and *ξ*_*N*_ are the corresponding drag coefficients. Plugging Eq. S19 into Eq. S17 yields Eq. S20.

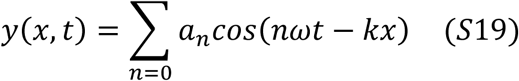

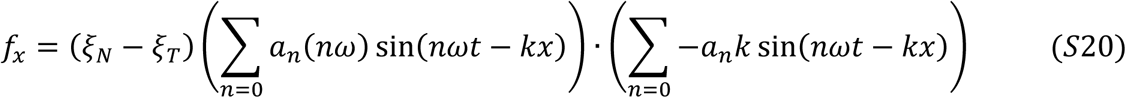

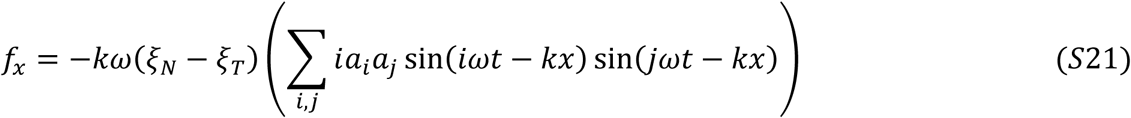

For *i* ≠ *j*, the time average of *f*_*x*_ would be zero, thus we only retain terms with *i* = *j*. This yields Eq. S22–23.

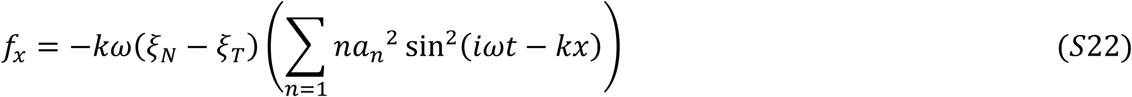

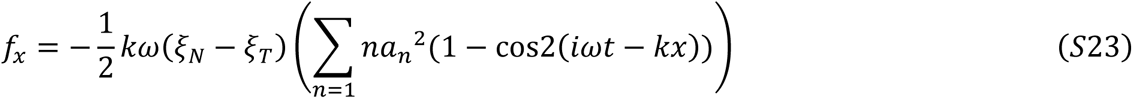

By calculating the time average of Eq. S23, the average force produced by each segment of the flagellum in the x direction can be described by Eq. S24.

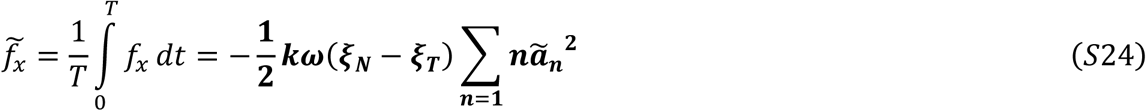

Integrating the 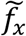 over the flagellum, the total force and velocity produced in the x direction is:

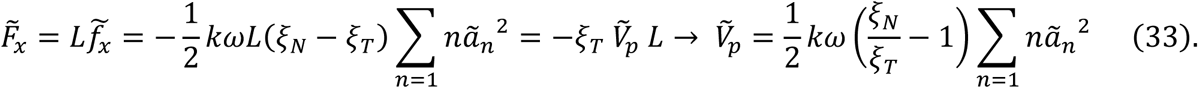

#### 2. Force in the y direction and the corresponding torque

Plugging Eq. S19 into Eq. S18 yields Eq. S25.

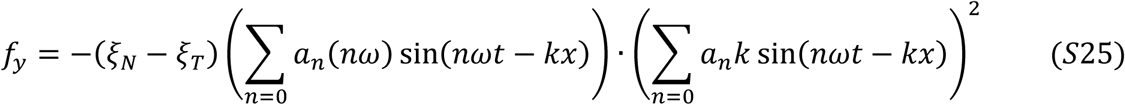

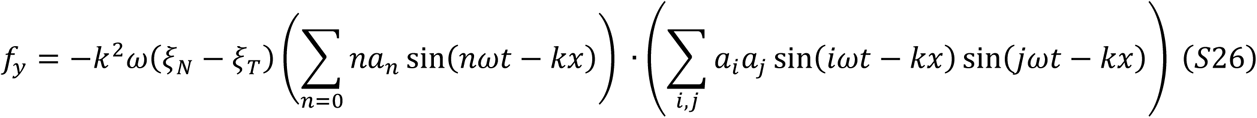

One can write out Eq. S26 in the form of Eq. S27.

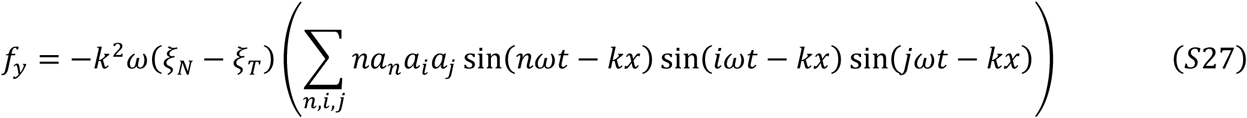

Since we are interested in time average values of *f*_*y*_, the following terms of Eq. S27 are non-zero:

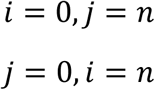

Therefore, Eq. S27 reduces to Eq. S28–29:

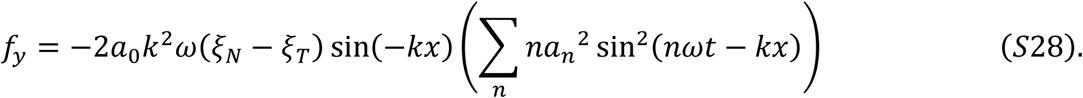

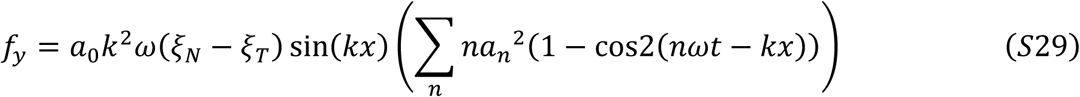

Taking the average of Eq. S28, the average force produced by each segment in the y direction is:

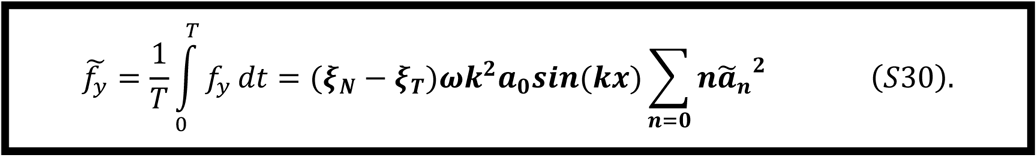

Interestingly, but not surprisingly, the force produced by each segment of the flagellum in the y direction is a function of x, meaning that the effect of the zeroth harmonic can be seen in the force produced in the y direction.

Integrating the forces produced by each segment in the y direction over the flagellum, the total force produced in the y direction becomes zero:

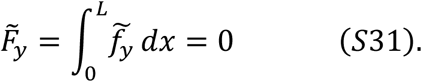

However, the total force produced in the front half of the sperm is non-zero and equal to the force produced in the rear half of the sperm:

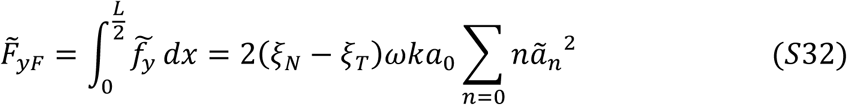

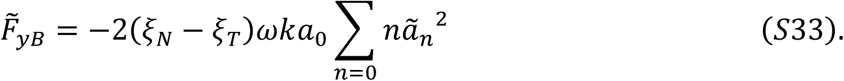

We calculated these two forces for the *γ* ratio, which is required for the Stokeslet description of the microswimmer model. Although the total force produced in the y direction is zero, the torque produced by the flagellum is not zero:

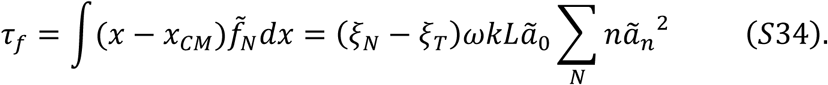

To find the angular velocity of the sperm 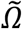, we need to calculate the torque produced by drag as well:

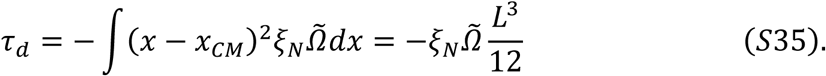

Considering the zero net-torque condition, we can simply find the angular velocity of the sperm:

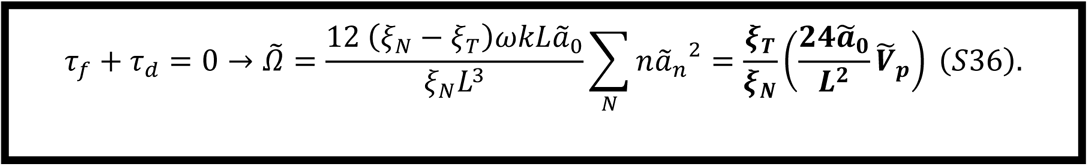

Since the curvature of the sperm trajectory is described by 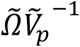 (in which 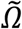 is the angular velocity of the sperm and 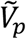 is the sperm velocity), the curvature can be defined as:

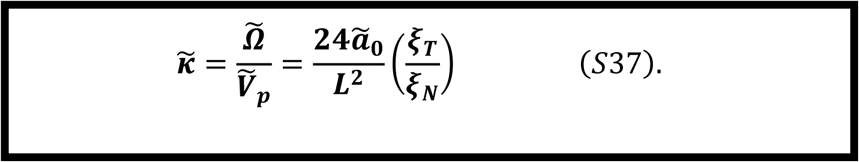

In turn, one can write out the *γ* ratio as:

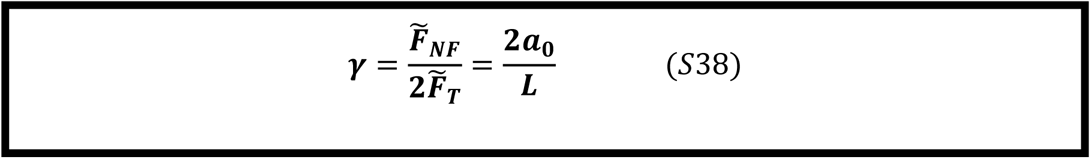

### IV. The analytic solution for the swimmer model using the Stokeslet description

The velocity field imposed by a source point is known as the Stokeslet (Eq. S39), i.e., the most fundamental solution for the Stokes equation. Based on the linearity in the Stokes equation, the contribution of the actively swimming sperm on the fluid flow can be described by superimposing the flow fields produced by each point force. For the sake of simplicity, we write out the imposed velocity field in three terms, including:

1. The velocity field imposed by 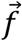 and 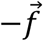 (Eq. S40),
2. 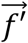 and 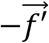 (Eq. S41), and
3. 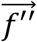 and 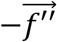 (Eq. S42). In which the magnitude of 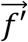 and 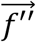 are equal to 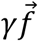.

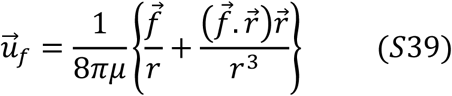

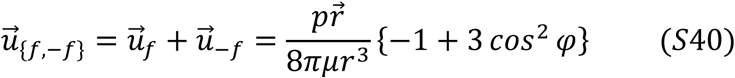

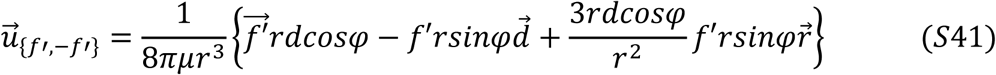

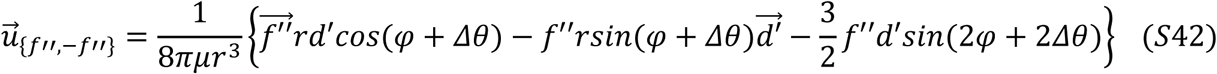

Using Eq. S40, S41, and S42, the velocity field imposed by the swimmer model is:

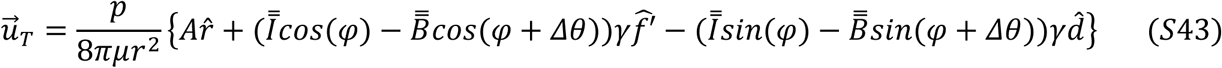

with

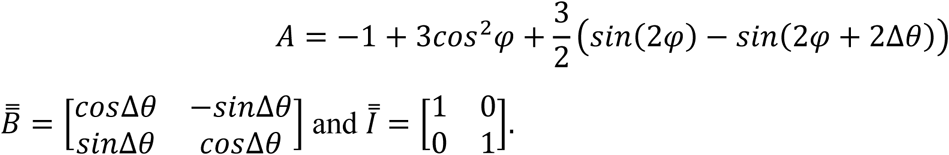

The influence of the no-slip boundary condition on the imposed flow is modeled by the mirror image of the swimmer in the boundary. Accordingly, the velocity field imposed on the sperm that causes the far-field attraction toward the wall is *u*_*T*_ with *r* = 2*h*, in which *h* is the distance between the sperm and the wall. For 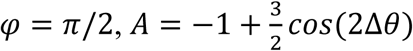,

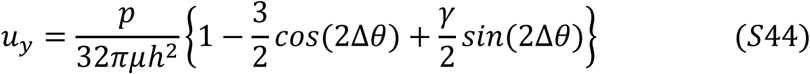

is the attractive velocity field imposed on the swimmer by the wall. This equation can also be written out as:

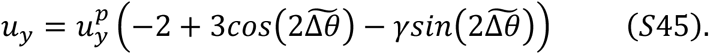

Interestingly, assuming *γ* ≪ 3, at 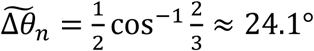, the swimmer experiences no attraction toward the walls and becomes neutral. Given the relation between 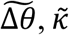, and 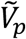, the corresponding curvature that is inversely related to the progressive velocity is:

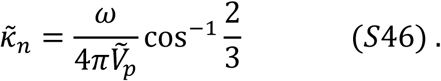

### V. The flow field produced by the progressive motility and circular motion

The flow field produced by the swimmer model in Fig. 4 is the superposition of the flow field produced by the components of progressive motility 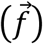 and circular motion (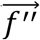 and 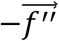) and their corresponding drags (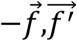 and 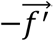). To identify the mechanism of reduction in the far-field attraction with the components of circular motion, we simulated the progressive term with corresponding drag (Fig. S4(A)) and the components of circular motion with corresponding drags (Fig. S4(B)), separately. Similar to Fig. 4 in the manuscript, the arrows show the normalized vector field while the magnitude of the flow is represented in color.

**Fig. S4.**
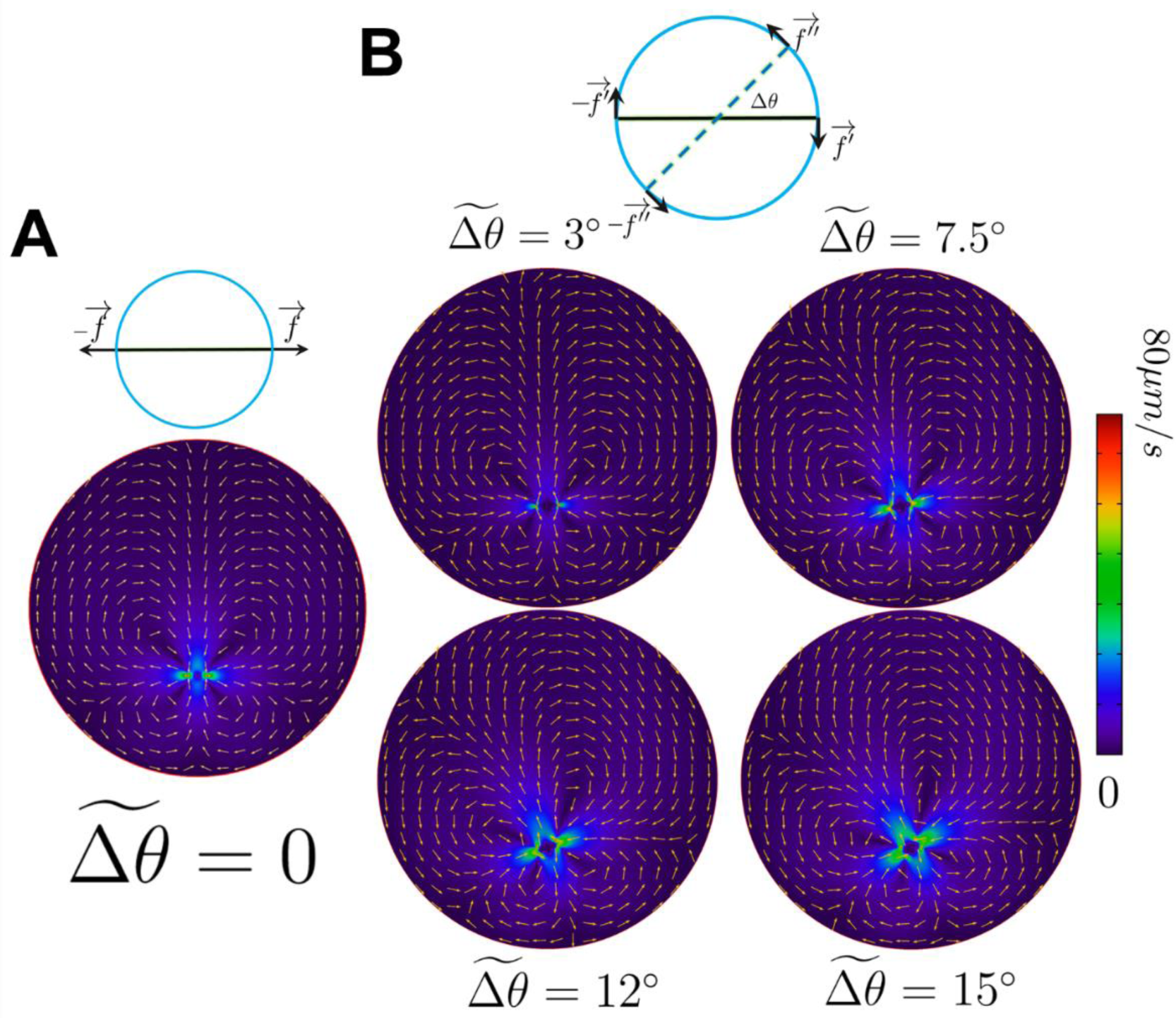
The flow field produced by the components of the progressive motility and the circular motion. (A) The flow produced by the progressive component of motility and its corresponding drag. (B) The flow produced by the components of circular motion and their corresponding drags, Δ*θ*, ranging from 3° to 15°. The arrows are the normalized direction while the colors represent the magnitude of the velocity field.

### VI. Lubrication approximation

The stress tensor is

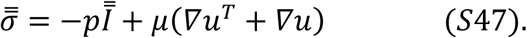

At distances adequately close to the wall, the contribution of the pressure dominates and 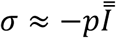. Accordingly, the torque exerted on the sperm by the boundary can be written out as

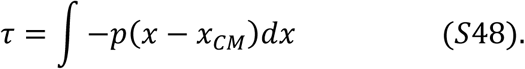

Neglecting the sperm mass, the net torque applied on the sperm is equal to zero, meaning that the drag torque cancels out the torque exerted by the wall. This constraint gives us

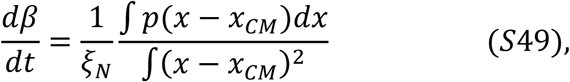

in which *β* is the angle of the sperm swimming direction with respect to the wall.

